# The fundamental principles of antibody repertoire architecture revealed by large-scale network analysis

**DOI:** 10.1101/124578

**Authors:** Enkelejda Miho, Victor Greifft, Rok Roškar, Sai T. Reddy

## Abstract

The antibody repertoire is a vast and diverse collection of B-cell receptors and antibodies that confer protection against a plethora of pathogens. The architecture of the antibody repertoire, defined by the network similarity landscape of its sequences, is unknown. Here, we established a novel high-performance computing platform to construct large-scale networks from high-throughput sequencing data (>100’000 unique antibodies), in order to uncover the architecture of antibody repertoires. We identified three fundamental principles of antibody repertoire architecture across B-cell development: reproducibility, robustness and redundancy. Reproducibility of network structure explains clonal expansion and selection. Robustness ensures a functional immune response even under extensive loss of clones (50%). Redundancy in mutational pathways suggests that there is a pre-programmed evolvability in antibody repertoires. Our analysis provides guidelines for a quantitative network analysis of antibody repertoires, which can be applied to other facets of adaptive immunity (e.g., T cell receptors), and may direct the construction of synthetic repertoires for biomedical applications.

## INTRODUCTION

The high diversity of antibody repertoires enables broad and protective humoral immunity, thus understanding their system sequence-related properties is essential to the development of new therapeutics and vaccines ^1,2^. The source of antibody diversity has long been identified to be the V-, (D- in the heavy chains) and J-gene somatic recombination ^3^. Further additions and deletions of nucleotides at the junctions of the gene segments generate a large collection of antibodies and B-cell receptors, which is called the antibody repertoire ^4,5^. Antibody identity (clonality) and antigen specificity are primarily encoded in the highly diverse junctional site of recombination in the variable heavy chain, known as the complementarity determining region 3 (CDR3) ^6^. Therefore, the similarity landscape of CDR3 sequences represents the clonal architecture of an antibody repertoire, which reflects the breadth of antigen-binding and correlates to humoral immune protection and function. However, due to limitations in technological sequencing depth and computing power, the fundamental principles that govern antibody repertoire architecture have remained unknown, thereby hindering a profound understanding of humoral immunity.

Recently, different aspects of network analysis have been employed to investigate antibody repertoires in health and disease. Antibody repertoire networks represent CDR3 sequence-nodes connected by similarity-edges ^7–11^. Sequence-based networks have first been used to show immune responses defined by similarity between clones, a proxy for clonal expansion ^8^. Network connectivity was later also used to discriminate between diverse repertoires of healthy individuals and clonally expanded repertoires from individuals with diseases such as chronic lymphocytic leukemia ^7^ and HIV-1 infection ^10^. A predominant part of network analysis has involved visualization of clusters and the display of clonal composition ^7–11^. Yet, visualization alone does not provide quantitative insights into the architecture of antibody repertoires and is limited to the informative graphical display of a few hundred nodal clones. It has been shown that the natural antibody repertoire exceeds the informative visualization threshold (hundreds clonal nodes) by at least three orders of magnitude (Glanville et al., 2009), a limit that previous research did not explore given the lower biological coverage (10^2^–10^3^ unique clones analyzed). Consequently, computational methods for constructing large-scale networks with more than 10^3^ nodes have remained underdeveloped in systems biology (Kidd et al., 2014). Furthermore, only networks expressing clonal similarity relations of one nucleotide (nt) or one amino acid (a.a.) between sequences have been analyzed so far, which is insufficient in covering the clonal relationships possible considering the extensive mutational landscape of somatic hypermutation ^12,13^. Thus, the lack of quantitative investigation of a relatively (and exceedingly) small subset of the antibody repertoire, with respect to clone numbers and network size, has limited the biological insight of repertoire architecture.

To reveal the fundamental principles of antibody repertoire architecture, we implemented a large-scale network analysis platform coupled to high-coverage antibody repertoire high-throughput sequencing data to answer the following questions: (i) Does sequence similarity among clones show reproducible signatures across individuals? (ii) How robust are antibody repertoires to removal of a fraction of clones, given their kinetics and rapid turnover? (iii) To what extent is the repertoire architecture intrinsically redundant? (Figure 1).

**Figure 1.**
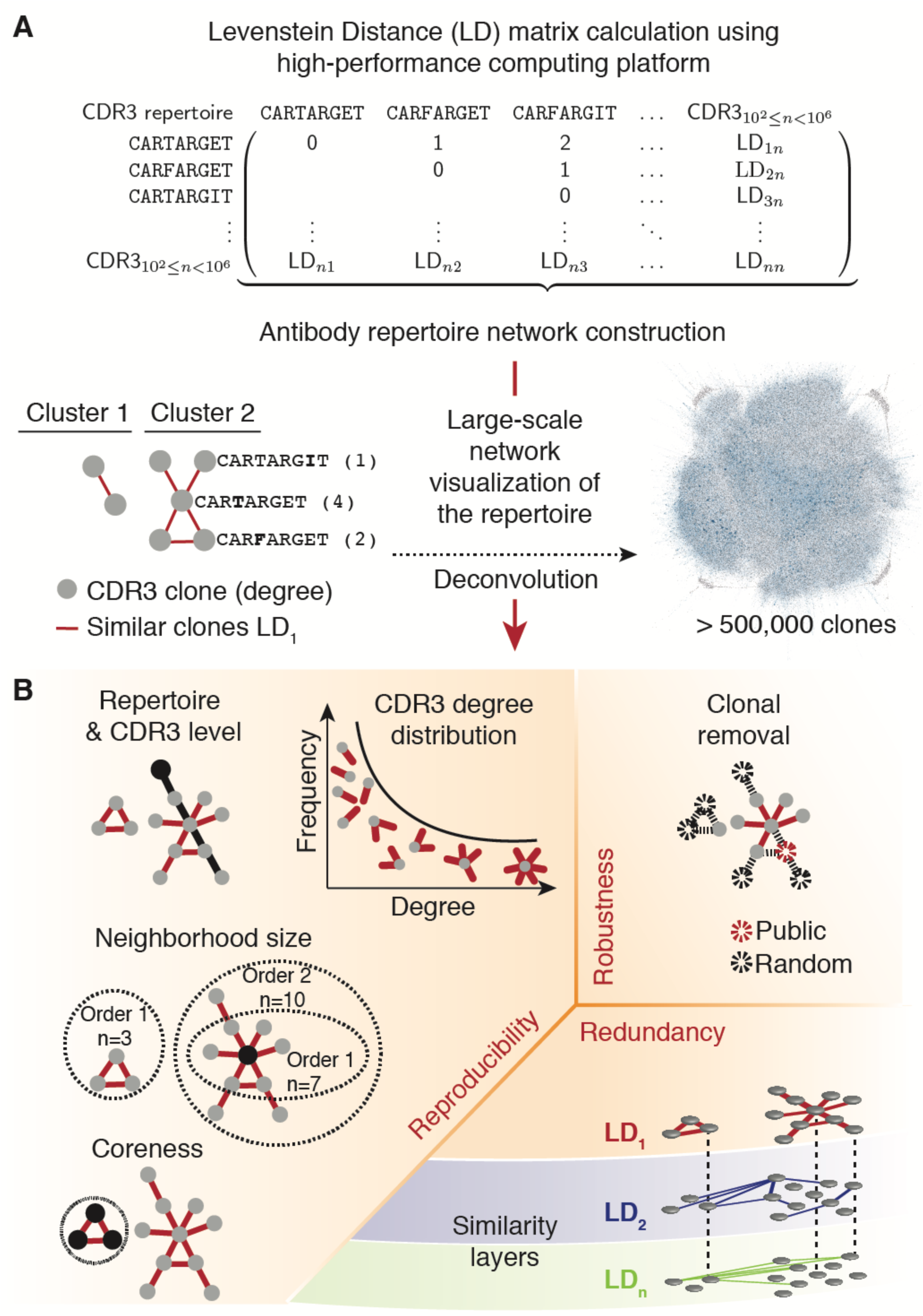
Large-scale network analysis reveals the architecture of antibody repertoires and its three main principles. A. Large-scale networks (>500,000 nodes) of antibody repertoires were constructed from the Levenshtein distance (LD, edit string distance) matrix of CDR3 clonal sequences (a.a) using a custom high-performance computing platform (see Methods). Networks represent antibody repertoires of similar CDR3 nodes connected by edges when amino acid CDR3 sequences differ by a predetermined LD. All clones of a repertoire connected at a given LD form a similarity layer (LDn).
B. Deconvolution of the complexity of antibody repertoire architecture was performed by quantifying (i) its reproducibility through global and clonal (local) properties, (ii) robustness to clonal deletion and (iii) redundancy across its similarity layers in the sequence space (Figure S1).

## RESULTS

### A high-performance computing platform for large-scale network analysis of antibody repertoires

The global landscape of antibody clonal similarities is vast and complex; for example, on the amino acid level, the size of the distance matrix of all-against-all sequences is ≈10^10^ for a representative repertoire of ≈10^5^ clones (murine naïve B cells). In order to extract the construction principles of antibody repertoires from the high-dimensional similarity space, we developed a large-scale network analysis approach, which was based on representing CDR3 a. a. clones as sequence-nodes connected by similarity-edges. Specifically, we developed a computational platform that leverages the power of distributed cluster computing, which is able to compute the extremely large distance matrices required for entire repertoires (≥10^5^ CDR3 sequences, Figure S1). Analysis of circa 10^5^ or more sequences is an intractable problem without parallel computing, thus our implementation utilized the Apache Spark ^14^ distributed computing framework to partition the work among a cluster of machines (Figure S1B). The construction of large-scale networks is computationally demanding: a large network comprised of 1.6 million nodes (simulated strings) required 15 minutes if the calculation was performed simultaneously on 625 computational cores (Figure S1C). Computational costs could have been lowered by performing network analysis on a subsample of the repertoire (e.g., 10^3^), as was done in previous studies ^7–11^. However, extensive analysis into sub-networks has revealed that they are not statistically representative of entire networks, specifically in that sub-network measurements are not always representative of key parameters such as degree distribution, betweenness, assortativity and clustering ^15,16^. Thus, it was imperative to construct and analyze large-scale networks based on a similarity distance matrix that covers the full clonal diversity of biological antibody repertoires.

Comprehensive biological sampling of antibody repertoires was ensured by the usage of high-throughput RNA sequencing data (≈400 million full-length antibody sequence reads) from murine B-cell populations, isolated at key stages in humoral development (data was provided by Greiff et al.). Data was analyzed from pre-B cells (pBC), naïve B cells (nBC) and memory plasma cells (PC) isolated from 19 mice, which were stratified into one unimmunized and three immunized cohorts. The experimental design allowed for the assessment of antibody architecture across several important parameters: i) across key stages of B-cell development, ii) before (pBC, nBC) and after antigen-driven clonal selection and expansion (PC), (iii) differences in the complexity of the protein antigen [hepatitis B surface antigen (HBsAg), ovalbumin (OVA) and nitrophenylacetyl-conjugated hen egg lysozyme (NP-HEL)], and (iv) across a scale of different repertoire sizes (10^2^–10^5^ of unique CDR3 clones). The experimental data provided maximal technological and high biological coverage (Greiff et al.), enabling comprehensive assessment of clonal diversity and the global similarity landscape and architecture of antibody repertoires.

For each sample (n=57, from 19 mice and three B-cell stages), antibody repertoire architecture was based on the pairwise a.a. sequence similarity of all clones (Levenshtein distance (LD) matrix, hereafter referred to as similarity layer, Figure 1A). When two sequences were similar within a defined threshold, they were connected in the repertoire network (i.e., similarities of 1 a. a. differences were captured in similarity layer 1, LD_1_, 2 a.a. in LD_2_ and so on).

### Global patterns of antibody repertoire networks are reproducible

In order to quantify the extent to which architectural patterns are reproducible across antibody repertoires, we analyzed the base similarity layer in antibody repertoires (similarity layer LD_1_). The base layer of the network organization provides information regarding the minimal differences (e.g., 1 a.a.) of all antibody sequences that compose the repertoire. Solely the base similarity layer, LD_1_, has previously been analyzed to describe antibody repertoires as networks ^7–11^. Although repertoires varied highly among mice (74–85% clonal variability, Figure S2A), we found a remarkable cross-mouse consistency in patterns of clonal interconnectedness (similarity of antibody sequences) within each B-cell stage: the average number of edges among clones (*E*_pBC_ = 230,395±23,048; *E*_nBC_ = 1,016,928±67,080; *E_PC_* = 45±10), the average size of the largest component (*pBC* = 46±0.7%; *nBC* = 58±0.5%; *PC* = 10±1.6%; Figure 2A) and cluster composition (Figure S2B) varied negligibly across mice (see Methods, Network analysis). Thus, although unique sequence composition varied substantially between individuals, the overall structure of the network remained similar.

**Figure 2.**
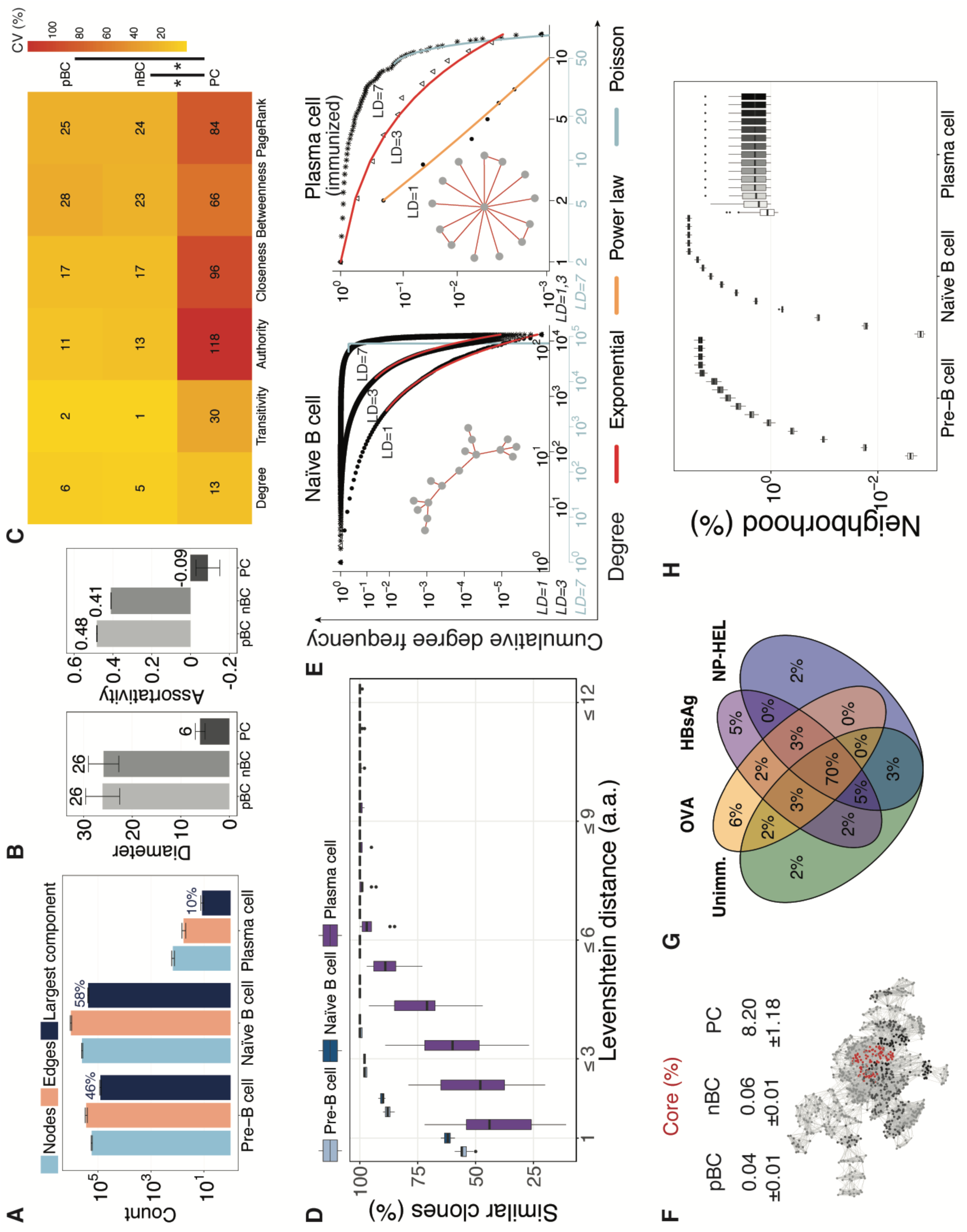
Global and clonal properties of antibody repertoire networks are reproducible. A. Network size of antibody repertoires. The y-axis indicates the absolute number count of CDR3 nodes, CDR3 edges (similarities) and CDR3 clones in the largest component. The mean percentage of the CDR3s belonging to the largest component by B-cell development stage is shown on top of the dark blue bar.
B. Global properties, diameter and assortativity coefficient are shown for pre-B cells (pBC), naïve B cells (nBC) and plasma cells (PC).
C. The mean value of the coefficient of variation for clonal properties in pBC, nBC and PC repertoires. Wilcoxon test, p_pBC,nBC/PC_ < 0.05 (see Methods).
D. Percentage of clones connected to at least one other clone in the repertoire at LD_1_, LD≤_2_,…, LD≤_12_ in pre-B cells, naïve B cells, plasma cells and randomly constructed CDR3 strings.
E. The power-law (orange), exponential (red) and Poisson (grey) distributions were fit to the cumulative degree distributions of naïve B cell and plasma cell (unimmunized) repertoires of a mouse for similarity layers LD_1,3,7_ (log-log scale). Representative clusters are shown for LD_1_.
F. Percentage of CDR3 clones (mean±s.e.m) that compose the maximal core. Subgraph of the maximal k-core (red), and k-1 (black), k-2 (dark grey) and k-3 (light grey) cores in a representative pBC sample (Unimmunized mouse n. 2).
G. Percentage overlap of CDR3 germline V-genes in the maximal core of nBC repertoires (n = 5 mice and data sets for Unimm, OVA, NP-HEL, n = 4 mice sets for HBsAg).
H. Normalized neighborhood size for orders n={1–10, 15, 20, 30, 40, 50} across CDR3 clones (similarity layer LD_1_). For 2A, B, D, barplots show mean±s.e.m; for 2A-E, each B-cell stage n = 19 mice.

Along B-cell development, PC repertoires were five-fold more disconnected than pBC and nBC networks (PC largest component was nearly 5 times smaller than pBC and nBC, Figure 2A), and their centrality was concentrated on specific clones compared to the homogeneously connected clones in pBC and nBC networks (centralization *z_PC_* = 0.05, density *D_PC_* = 0.01, *z_pBC,nBC_* ≈ *D_nPC,nBC_* ≈ 0, Figure S2C). Compared to pBC networks, nBC were on average 4–5 times larger and showed a higher average degree (*k*_pBC_ = *3*, *k*_nBC_ = 5, *k*_PC_ = 1, Figure S2B) although both pre- and naïve B-cell repertoires had identical diameter (*d*_pBC,nBC_ = 26, *d*_PC_ = 6, Figure 2B), indicating a similar coverage of the sequence space. We observed that clones in pBC and nBC repertoires connected to comparable clones in terms of degree (assortativity ^17–19^, *r*_pBC_ = 0.48, *r*_nBC_ = 0.41), whereas PC networks were consistently disassortative: their highly connected clones were linked to clones with few connections (r_PC_ = −0.09, Figure 2B). The characterization of the global patterns of antibody repertoire networks indicated that pBC, nBC and PC repertoires were reproducible. pBC and nBC clones cover a larger diversity space than clones in PC repertoires, where sequence similarity showed to be centralized and targeted towards certain clones.

### Clonal features of antibody repertoire networks are reproducible

Antibody repertoire architecture was also reproducible at the level of clonal (local) features in pBC and nBC networks, which were characterized by a low variability (coefficient of variation, CV) across various clonal parameters. The low variability of clonal parameters in pBC and nBC networks (*CV*_pBC_ = 2 – 28%, *CV*_nBC_ = 1 – 24%) was in contrast to the higher variability observed in PC repertoires (*CV*_PC_ = 13 – 118%, Figure 2C). Specifically, low variability across different individuals was observed in several average clonal parameters such as degree, transitivity, authority and PageRank, closeness and betweenness. Variation analysis of the similarity degree indicated that the average number of similar clones to each of the clones in a repertoire varied marginally in pBC and nBC (*CV*_pBC,nBC_ = 5,6%). Transitivity showed that the similarity between clones both similar to a third CDR3 clone varied only negligibly between individuals (*CV*_pBC,nBC_ = 1,2%). Authority and PageRank showed that the centrality of a CDR3 in the repertoire topology varied respectively *CV_pBC,nBC_* = 11% and 25% across individuals, suggesting that individual repertoires were centered variably around certain CDR3 clones which were centers of highly connected (similar) clonal regions compared to less connected regions in the same repertoire network.

Closeness analysis revealed that an analogous number of similarity edges were required to access every other CDR3 from a given CDR3 clone in antibody repertoire networks of different individuals, as the similarity of a clone to every other CDR3 clone in the repertoire varied by *CV*_pBC,nBC_ = 17%. Betweenness, the “bridge” function of a clone in sequence similarity, varied slightly across individuals with *CV*_pBC,nBC_ = 28%, suggesting a comparable structure of the similarity route function of CDR3 sequences in these repertoires. These characteristics reflect the transversal diversity of pBC and nBC antibody repertoires where the clones cover a larger space and their similarity is more homogenously distributed at the global repertoire level.

Although a higher variability was detected across PC repertoire networks (Figure 2C), clonal parameters were specific to B-cell stages (p_pBC,nBC/PC_ < 0.05): PC clones possessed higher centrality compared to pBC and nBC (closeness^20^, eigenvector^21^, and PageRank^22^), while antigen-inexperienced clones bridged sequence similarity (betweenness ^23^, Figure S2D). Furthermore, in contrast to pBC and nBC, PC network clonal parameters correlated with CDR3 frequency (clonal degree median *r*_Pearson_ = 0.55, betweenness *r*_Pearson_ = 0.82) suggesting that clonally expanded CDR3 sequences were structural centers of similar clones (Figure S2E, F). CDR3 authority correlated positively with germline V-gene frequency in PC clones (*r*_Pearson_ =0.39), denoting the potential role of the V-gene usage in the centralization of these networks (Figure S2G). Thus, certain high frequency V-genes predispose clones to be highly connected and similar to other clones.

### The structure of antibody repertoires is reproducible and depends on the immune status

Network analysis revealed that antibody repertoires were constricted along B-cell development throughout all similarity layers. At LD_1_, 44–62% of clones were similar (connected) to at least one other clone in all B-cell stages, revealing a high degeneracy in clonal generation and selection (Figure 2D). This indicated that nearly half of antibodies in the respective repertoires had similar clones, thus demonstrating the extent of constriction present in antibody repertoires.

In order to understand if such degeneracy in CDR3 sequence similarity translated into reproducible repertoire network structures ^21^, we determined the clonal empirical degree distribution. The degree distribution is a distinctive feature of different types of networks and it provides an immediate indication of how similarities (degrees) between antibody sequences are distributed in repertoires. Analysis of the cumulative degree distribution revealed that antigen-inexperienced pBC, nBC and unimmunized PC repertoires were exponentially distributed (LD_1_), whereas PC repertoires of immunized cohorts were power-law distributed (base similarity layer LD_1_, Figure 2E, Figure S3D, E, F, G). Clusters of connected CDR3 clones showed a typical tree-like structure for pBC and nBC, and a star-like structure for PC. The structure of the network suggested an extended and chain-wise sequence similarity of the antibody clones in pBC and nBC repertoires and targeted expansion of certain clones in PC after immunization.

In order to prove the tree-/star-like hypothesis and further investigate the sequence similarity space, we performed *k*-core ^24^ decomposition and neighborhood analysis (Figure 2F, G, H). The *k*-core decomposition revealed that the largest *k*-cores (after all external shells with *k*<*k*_max_ were removed, where *k* is the degree, i.e. number of similar clones, see Methods) of pBC and nBC (0.04% and 0.06% of CDR3 clones in *k*-core, respectively) were 200-fold smaller than those of PC (8.2%, Figure 2F). Antigen-inexperienced repertoires were thus characterized by larger coreness values (>20), signifying a more layered structure of CDR3 similarity (Figure S2J, K) and confirming their tree-like structure. Furthermore, the high convergence of V-genes at the core-level of antibody repertoire networks (pBC=50%, nBC=70%, PC=1–10%, Figure 2G), in contrast with the low exact CDR3 sequence core-overlap (Figure S3A, B, C), suggested a genetically determined origin of the structure.

The average CDR3 neighborhood size, which designated the set of similar CDR3 clones along each sequential step of similarity from a certain clone (orders n=1–50), was order-independent in PC and plateaued at 2% of the network, confirming that PC clones were connected to one central clone in a star-like similarity structure, reflecting clonal selection and expansion signatures. Neighborhood size ^25^, the number of similar clones to each clone, increased order-wise in antigen-inexperienced cells up to 34% (Figure 2H), signifying tree-like similarity structures that enable maximal exploration of sequence space within the genetically predetermined repertoire constriction space, suggesting that antibody repertoires are evolutionarily wired to respond to diverse antigenic stimuli.

### Antibody repertoires are highly robust systems

We hypothesized that the reproducible architecture of antibody repertoires may have evolved to be robust to fluctuations in clonal composition. It is known that antibody repertoires are very dynamic systems characterized by a high turnover rate ^26–28^. Therefore, we investigated the robustness of antibody repertoire architecture to clonal removal (deletion).

It is has been recently established that individual repertoires have public clones, which are defined as identical clones present in multiple individuals ^29^. While mostly distinct, antibody repertoires possessed a fraction of public clones (15–26% along B-cell development, Figure S2A). Given their regular presence, we determined if public clones were essential to the maintenance of antibody repertoire architecture. We found that the highest authority clones were public (Figure 3A) and up to 74% of private clones (specific to an individual) were connected to at least one public clone (Figure S2I). To quantify the extent to which public clones maintain the architecture of antibody repertoires, we tested the effect of removing public clones on CDR3 degree distributions. In pBC and nBC, removal of all public clones transformed their network structure from exponential to power-law; in contrast, removal of public clones led to no change in PC network structure (Figure 3B). To assess if such a structural shift was specifically due to the deletion of public clones, we removed (repeatedly) random subsets of clones representing a similar fraction of public clones. The structure of antibody repertoires was robust along B-cell stages at up to 50% removal of random clones. The same structural shift in repertoire structure caused by the deletion of public clones could only be replicated by removing 90% of random clones (Figure 3C). Therefore, public clones represent pillars that are critical for maintaining the architecture of an antibody repertoire, and the robustness of this architecture suggests a functional immunity is preserved even after extensive (random) loss of antibody clones (or B cells).

**Figure 3.**
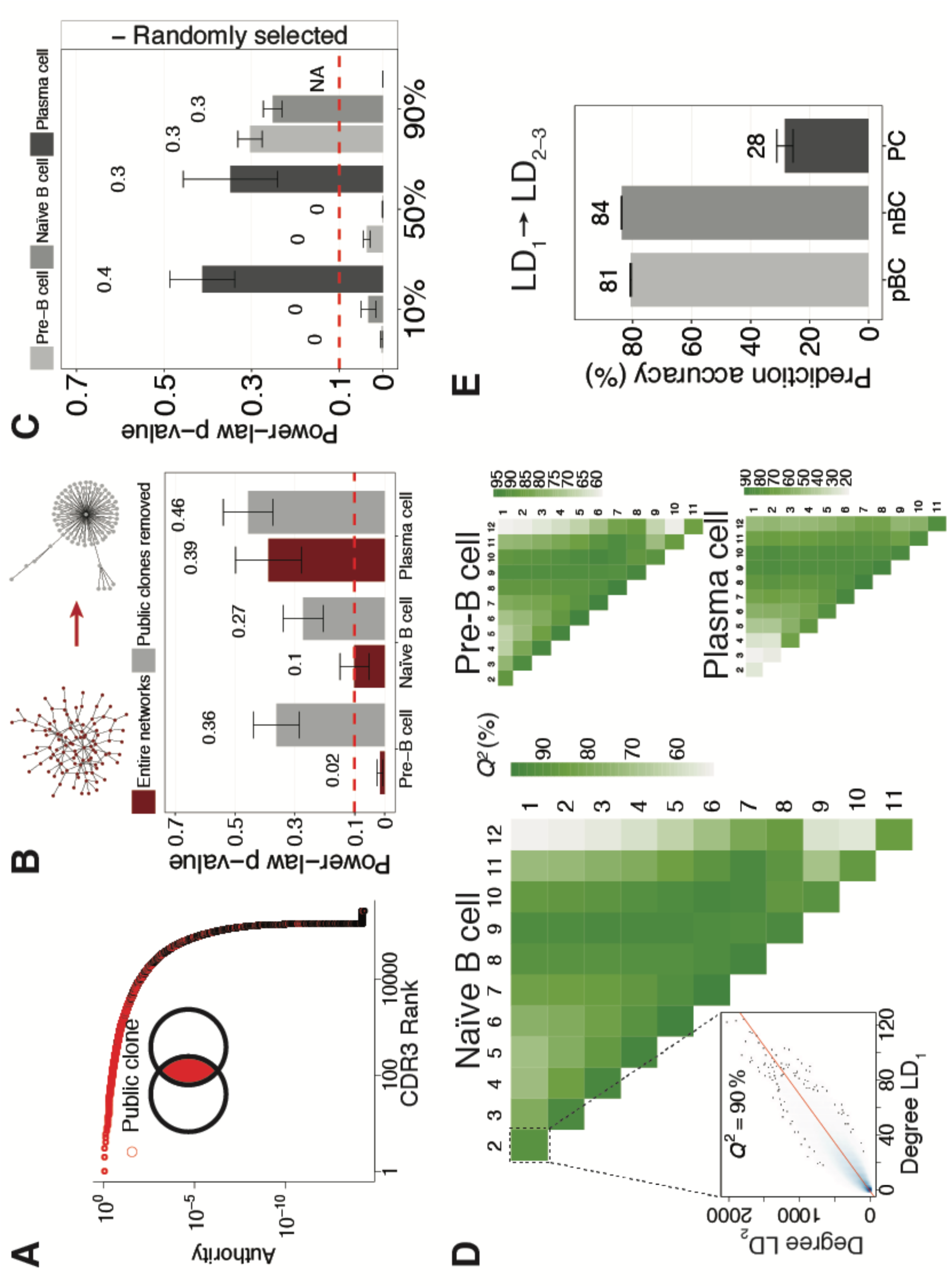
The architecture of antibody repertoires is robust and redundant. A. CDR3 clones of an exemplary naïve B-cell repertoire (from OVA-immunized mouse n. 1) have been ordered from increasing to decreasing frequency (CDR3 rank). Public clones are color-coded in red.
B. Bootstrapped p-values of the power-law fit are shown for complete antibody repertoires and after removing public clones. Power law is a good fit to degree distributions for p-values above the dashed red line (p-value = 0.1). Examples of exponential (red) and power-law (grey) networks are shown on the top panel.
C. CDR3 clones were removed randomly at 10%, 50% and 90% from each original repertoire (20 times) and the power-law distribution was fit to the cumulative degree distributions of the remaining CDR3 clones. A p-value=0.1 is indicated as a red dashed line. In PC samples a fit was not feasible after removal of 90% of CDR3 clones (NA).
D. Heatmaps indicate the mean prediction accuracy (*Q^2^*, leave-one-out cross-validated *R^2^*) of similarity layer LD_1_ versus similarity layers LD_2–12_. The scatterplot shows *Q^2^* for LD_1_ vs. LD_2_ for each CDR3 clone.
E. Prediction accuracy (*Q^2^*) for LD_1_ vs. LD_2_ and LD_3_. For 2B, C, E, barplots show mean±s.e.m.

### Antibody repertoires are evolutionary redundant

Redundancy is a hallmark of robust systems; for example, redundancy in genes with the same function is the main mechanism of robustness against mutations in genetic networks ^30^. To investigate the extent of redundancy within antibody networks, we examined whether their architecture at the base similarity layer (LD_1_) was manifested in higher order similarity layers (LD_>1_). Differences greater than one a.a. between antibody sequences could represent the potential personal scenarios of antibody repertoire evolution (somatic hypermutation ^12,13^), a result of successful survival through selective processes. Specifically, if a clone connected to many other clones in the LD_1_ similarity layer mutates into a similar clone at a specific a.a. position, this *potential* clone will be connected to many clones in the LD_2_ similarity layer. Thus, higher order similarity layers can serve as surrogates for the evolution of potential antibody repertoires from antigen-inexperienced B-cell populations.

To quantify the extent of redundancy across similarity layers, we calculated the prediction accuracy of LD_1_ versus similarity layers LD_2–12_ using a leave-one-out cross-validation approach (Figure 3D, Figure S3H and S3I). Specifically, quantitative redundancy was low in PC (LD_1_→LD_2–3_ prediction accuracy was 28% on average); however, LD_1_ of pBC and nBC predicted CDR3 degree profiles of proximal similarity layers LD_2–3_ with ≥80% accuracy (Figure 3D and 3E), thereby indicating a high redundancy in antibody repertoire architecture. This high redundancy is explained by the structure of the antibody networks (Figure 2E–H). Although the distance between proximal similarity layers (LD_1_ to LD_3_) seems small (1–3 a.a. CDR3 sequence differences), it represents ≈20% of potential change in clonal a.a. sequence (99% of CDR3 clones are 4–20 a.a. long), which is in the range of highly mutated antibodies (e.g., broadly-neutralizing HIV-specific ^31^). Therefore, redundancy in the antigen-inexperienced repertoire is maintained throughout a large sequence space and provides details on the pre-programmed evolvability ^32,33^ of antibody responses.

## DISCUSSION

In summary, leveraging a custom-developed analysis platform for generating large-scale networks from datasets of millions of unique sequences, we have discovered three fundamental principles of antibody repertoire architecture: (i) reproducibility (ii) robustness and (iii) redundancy. We were able to detect a high cross-individual reproducibility by quantifying network parameters ^17–19^ at the global (size, diameter and assortativity) and clonal level (degree, transitivity, authority, closeness, betweenness, PageRank) of antibody repertoires along B-cell development. Importantly, the reproducible clonal similarity properties were suggestive of the underlying immunobiology of each B-cell stage: antigen-inexperienced repertoires covered an extended sequence diversity space (tree-like exponential similarity structure) to counter high antigen diversity whereas, antigen-experienced repertoires presented a centralized network structure (star-like, power-law), with many clones being similar to one central clone possibly originating from antigen-dependent clonal expansion and selection ^34^.

Large-scale network analysis of entire antibody repertoires revealed that these systems are robust enough to be amenable to subsampling, which is in contrast to other systems (Lee et al., 2006; Sethu and Chu, 2012). Specifically, we showed that the structure of antibody repertoire networks was robust to extensive subsampling of up to 50% of the clones. This result is crucial for the network analysis of human antibody repertoires, where biological subsampling remains an important problem ^35,36^. The robustness of antibody repertoires might also explain their functionality despite the large fluctuations of antibody repertoire composition over time ^26–28^. Interestingly, the structure of antibody repertoires was fragile to the removal of public clones. The crucial role that public clones play as pillars of antibody repertoire architecture was revealed by large-scale networks, yet future research will need to determine the functional role (antigen specificity) of public clones in a humoral response.

We found that antibody repertoires presented intrinsic redundancy across similarity layers. This means that not only minimal differences (1 a.a. of the base layer LD_1_) but also further diversification (> 1 a.a. differences between antibody sequences) may be hardcoded into the constricted sequence space of antibody repertoires, thus rendering their evolvability robust (analogously to other biological systems such as transcription factor networks ^33^).

This work delineates guidelines for the large-scale network construction and analysis of large and diverse immune repertoires. In particular, our network analysis approach can be used where a partial biological coverage of the repertoire is available, although this might depend on the B-cell stage, species, and similarity layer investigated. The network quantitative analysis of global and clonal properties of adaptive immune repertoires (antibody and T cell receptor repertoires) in health and disease is essential to comprehensively understand their architecture and may resolve limitations arising from visualization of graphics featuring high-dimensional data.

The three fundamental principles of the architecture of antibody repertoires uncovered here through network analysis may serve as a blueprint for the construction of synthetic antibody repertoires, which may be used to simulate natural humoral immunity for monoclonal antibody drug discovery and vaccine development ^32,37^. Large-scale antibody network analysis could be useful in personalized medicine in the prediction of immunity scenarios for repertoire-transforming diseases such as autoimmunity or lymphomas, which lead to major alterations in repertoire composition ^38,39^; this may allow for interventions to modify disease progression on the repertoire level by precision therapeutic clonal targeting. Finally, we believe the stage is set for a rapid progression of the present guidelines into what was long ago envisioned by Niels K. Jerne ^40^: the field of *network systems immunology*, which offers the potential to obtain greater understanding of the complexity of immune responses.

## METHODS

### EXPERIMENTAL MODEL AND SUBJECT DETAILS

#### Dataset

The dataset analyzed was produced as described by Greiff et al. in the attached manuscript. Briefly, murine B-cell populations of pre-B cells (pBC, IgM, bone marrow), naïve follicular B cells (nBC, IgM, spleen), and memory plasma cells (PC, IgG, bone marrow) were sorted using fluorescence-activated cell sorting (FACS) from C57BL/6J mice unimmunized (n=5) or prime-boost immunized with alum-precipitated antigens: nitrophenylacetyl-conjugated hen egg lysozyme (NP-HEL, n=5), ovalbumin (OVA, n=5) or Hepatitis B virus surface antigen (HBsAg, n=4). Following total RNA extraction, full-length antibody variable heavy chain (VDJ) libraries were generated by a two-step PCR process, as described previously ^41^. Libraries were sequenced using the Illumina MiSeq (2x300bp) platform. Mean Phred-scores of raw data were ≥30. Approximate paired-end reads (full-length VDJ) were: pBC 5×10^6^ reads, nBC 10×10^6^ reads and PC 4×10^6^ reads.

## METHOD DETAILS

### Data preprocessing and CDR3 clonal analysis

Antibody sequences have been preprocessed and VDJ annotated with MiXCR ^42^ and further filtered to retain only those sequences that had CDR3 length ≥ 4 a.a. and occurred more than once in each CDR3 repertoire data set (Figure S1A). Clones were defined by 100% a.a. sequence identity of CDR3 regions. CDR3 regions were defined by MiXCR according to the nomenclature of the Immunogenetics Database (IMGT) ^43^.

#### Network construction

To construct networks (graphs), a sparse triangle matrix of pairwise Levenshtein distances (LD) between CDR3s must first be computed. For small samples (up to 100,000 unique CDR3 sequences) such a calculation is relatively quick on a single computer. However, due to the N^2^ complexity of required calculations, computing the pairwise matrix for samples of >100,000 unique CDR3 sequences becomes prohibitively expensive. To perform these computations, we developed software that utilizes the Apache Spark (2) distributed computing framework to partition the work among a cluster of many machines (Figure S1B). We chose specifically Apache Spark because i) its deployment is very flexible with regard to underlying computing infrastructure and ii) for similarity layers LD_>1_, the networks become extremely large and difficult to process. In these cases, our package can take advantage of the Spark GraphFrames distributed graph library ^44^, which allows scaling to even larger samples with millions of sequences (Figure S1C). With this approach we were able to compute the distance matrices for large samples (>100,000 unique CDR3 sequences) within minutes (Figure S1, B and C). In addition to the computational complexity inherent in creating the distance matrix, the construction of networks for large LD is very computationally and time-wise costly. We therefore avoided constructing networks altogether for calculating the node degrees and instead used a map-reduce distributed algorithm. For practical purposes, the construction of small networks was performed using the Networkx library ^45^. For generating and outputting the largest graphs to disk in common network formats, we used the efficient graph-tool library (https://graph-tool.skewed.de/, ^46^). For manipulating and analyzing the largest networks, our software package took advantage of the Spark GraphFrames distributed graph library ^44^.

The software was developed in python (https://www.python.org/) using the Numpy/Scipy ^47^ scientific libraries for matrix and array manipulation and Apache Spark ^14^ as the distributed backend. Our software package for antibody repertoires *imNet* is available (https://github.com/rokroskar/imnet) and includes tutorials and demos, including scripts to set up the distributed computation environment on commonly-used compute cluster infrastructure. The results shown in this work were obtained using 1–625 cores of the *Euler* parallel-computing cluster operated by ETH Zürich.

## QUANTIFICATION AND STATISTICAL ANALYSIS

### Degree distribution fits

Degrees (number of similar CDR3 sequences to a specific CDR3 sequence) were calculated for each of the similarity layers LD_1–12_ for each CDR3 sequence in each sample. CDR3 with zero degrees that were not similar to any other CDR3 in the network were excluded in order to fit degree distributions. The power-law, exponential and Poisson distributions were fitted to the empirical degree distributions of the networks, constructed as described in *Network construction*, by estimating x_min_ (estimated lower degree threshold by minimizing the Kolmogorov-Smirnoff statistic ^48^) and optimizing model parameters using the poweRlaw ^49^ package. We first discriminated if the power-law distribution could describe the best fit to the degree distribution by bootstrapping 100 times the power-law p-value obtained from each sample after estimating x_min_. Following the approach described by Virkar and Clauset ^50^, a p-value ≥ 0.1 indicated that the power-law distribution described the degree distribution (Figure S1A). To determine the degree distribution in cases where the power law was not the best distribution fit (p-value < 0.1), we compared the exponential and the Poisson fits. Two-sided p-value≈0 indicated that the fitted models could be discriminated, and one-sided p-value≈1 indicated that the first (for example exponential) model was the best fit for the data ^49^.

### Robustness of the architecture of antibody repertoire networks

Public clones were defined as clones shared among subjects in a cohort (Figure S2). In order to assess the robustness of the architecture of antibody repertoire networks we removed public clones from each sample-repertoire. As controls, we performed repeated removal (20 times) of randomly selected clones in the size of public clones. The p-values for the power-law fit were calculated after 100x bootstrapping for each repertoire; one-sided and two-sided p-values were used for the comparison between the exponential and the Poisson fits (see *Degree distribution fits*).

### Network analysis

Drawing from network theory ^51^, we translated the concepts of network analysis ^18^ to antibody repertoires. An antibody repertoire network is an undirected *graph* G = (V, E) described as a set of *nodes* (CDR3 vertices, V) together with a set of connections (similarity edges, E), representing the adjacency matrix *A* of pairwise Levenshtein distances (LD) between CDR3 a.a. sequences 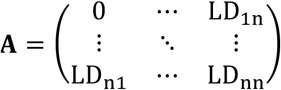.

In the context of antibody repertoires, we let N = |V| and L = |E|. The *order* of a graph N represents the number of its unique CDR3 clones (nodes). The *size* of a graph L is the number of its CDR3 similarity connections (edges). The degree k, that represents the edges connected to a node, describes the count of all similar CDR3 clones to a CDR3 based on LD. Because the *degree* indicates how active a node is, it could be interpreted as a measure of how central a CDR3 clone is in the antibody repertoire network. In simpler terms, it quantifies the number of CDR3 clones that are similar to a certain CDR3, and thus the potential development or the evolutionary routes to this CDR3.

The *average degree* 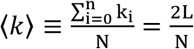 is the average number of similar CDR3 clones. The *degree distribution* = N_k_/N, defined as the fraction of nodes with degree *k* (N_k_) in total nodes, represents the fraction of CDR3 clones that have the same number of similar CDR3s. The *cumulative degree distribution* P_k_ 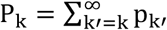 describes the fraction of nodes with degree greater than or equal to k’. In Erdős–Rényi (ER) random graph models, degrees follow a Poisson distribution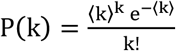 in the limit of large numbers of nodes, while degree distributions have an exponential tail 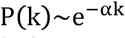in exponential networks ^52^.

Global characterization ^18^ described the network as a whole, such as degree distribution, centralization, largest component, diameter, clustering coefficient, assortativity and coreness. The *centralization* analysis indicates if the network is homogeneous (clones are connected in the same way) or is centered around certain nodes (highly connected clonal regions compared to less connected regions in the same network). The *largest component* is the largest cluster of connected CDR3 clones. The *diameter* (*d*) is the maximum distance (shortest path between two nodes) between any pair of CDR3 sequences. The *clustering coefficient* (*C*) represents the probability that neighbors of a node are also connected, which translates in antibody repertoires as the probability that CDR3 clones similar to a specific CDR3 are also similar among one another. Network *density* (*D*) is the ratio of the number of edges (CDR3 similarities) and the number of all possible edges in the network. The *assortativity coefficient* (*r*) indicates if nodes in a network connect to nodes with similar characteristics. It is positive if nodes tend to connect to nodes that are similar to them (i.e. highly connected CDR3 sequences are similar and connect to highly connected CDR3 sequences), and negative otherwise. *Coreness* is a measure of the network’s cohesion and allows one to understand the global network structure and is useful in comparing complex networks by analyzing the subsets of CDR3-cores that form layers in the antibody repertoire. *K*-core decomposition is a process that is performed by iteratively removing shells of all vertices of degree less than *k* (*k*<*k*_max_) leaving the *k*-cores of a network (its connected component). The *k*-core of a graph is the maximal subgraph in which each node has at least degree *k*. We have computed the maximal *k*-core of antibody repertoire networks (the innermost core, *k*_max_) and the core distribution along *k* degrees.

Clonal (local) characterization of antibody repertoires was performed by analyzing local properties of the networks ^18^. The importance of CDR3 clones was measured by calculating the authority ^53^, eigenvector ^22^ and PageRank ^23^ scores of each node in repertoire networks. In particular, the *authority* (*a*) of nodes is defined as the principal eigenvector of the transpose matrix t(**A**) * **A**, where **A** is the adjacency matrix of the network. Eigenvector centrality indicates the centrality of a CDR3 clone, not only dependent on the number of similar CDR3 (number of degree, connections) but also on the quality of those connections: CDR3-nodes with high eigenvector values are connected to many other nodes which are, in turn, connected to many others (and so on). *PageRank* measures the importance of the similarity between two CDR3 clones within the network extending beyond the approximation of a CDR3 importance or quality. *Closeness* (*centrality* ^21^) (*c*) was calculated to measure how many steps were required to access every other CDR3 from a given CDR3 clone in antibody repertoire networks. We calculated the normalized closeness by multiplying the raw closeness by n**-**1, where *n* was the number of nodes in the network. *Clique* analysis identified maximally-connected subgraphs (a subset of nodes) in which every CDR3 was similar to every other CDR3 sequence and the largest clique was the maximal completed subgraph which had more nodes than any other clique in the network. The node betweenness (b) is the number of geodesics (shortest paths) going through a node and indicates the “bridge” function of a CDR3 sequence. Network properties were calculated using the igraph ^54^ R package.

### Quantifying the predictive performance (Q2) of linear regression models

The predictive performance (*Q*^2^) of each linear regression model (***Y*** = ***X**β* + ***ε***) was calculated using leave-one-out cross-validation (LOOCV):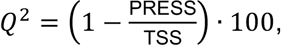 where PRESS is the predictive error sum of squares 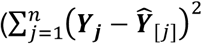 with 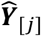 denoting the prediction of the model when the y-th case is deleted from the training set and TSS is the total sum of squares 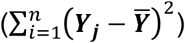 (Greiff et al., 2012). ***X*** and ***Y*** are CDR3 degree vectors of repertoires at each LD_1–12_. LOOCV was performed using the forecast R package ^55^. Cross-validation was used because, in contrast to regular regression analysis, it enables the quantification of the predictive performance of each regression model.

### Simulated networks

Networks (nodes V=10^2^–10^5^) were simulated with the ER, exponential and power-law models using base R ^56^ and igraph ^54^. Random networks were simulated according to the ER model, exponential networks were simulated setting a probability of a connection between two nodes p=0.5 and scale-free networks were simulated using the Barabási-Albert model (Barabási and Albert, 1999).

### Graphics

Graphic representations were produced using base R ^56^ and the ggplot2 R package ^57^. Heatmaps were produced using the NMF package ^58^. Networks and network clusters visualization were performed using igraph ^54^ employing the Fruchterman–Reingold force-directed and Kamada–Kawai layout algorithms. Large-scale networks (Figure 1a) were visualized using Gephi (version 0.9.1) ^59^; node size was scaled 10–100 proportional to the degree of a node and a blue to grey color gradient was applied to nodes from high to low degrees.

### Statistical significance

Statistical significance was tested using the Wilcoxon rank-sum test. Results were considered significant for p<0.05.

### DATA AND SOFTWARE AVAILABILITY

Antibody repertoire sequencing data analyzed is available with ArrayExpress accession number: E-MTAB-5349. Software is available at https://github.com/rokroskar/imnet.

## Acknowledgments

We thank Manuel Kohler from the Scientific IT Services of ETH Zürich for technical support. We thank Dr. Antonios Garas, Ulrike Menzel and Simon Friedensohn for scientific discussions, and Dr. Laura Prochazka, Alexander Yermanos and Cédric Weber for critically reading the manuscript. This work was funded by the Swiss National Science Foundation (Project no: 31003A_143869, 31003A_170110 to STR), SystemsX.ch – AntibodyX RTD project (to STR), Swiss Vaccine Research Institute (to STR). The professorship of STR is made possible by the generous endowment of the S. Leslie Misrock Foundation.

## SUPPLEMENTARY INFORMATION

**Figure S1.**
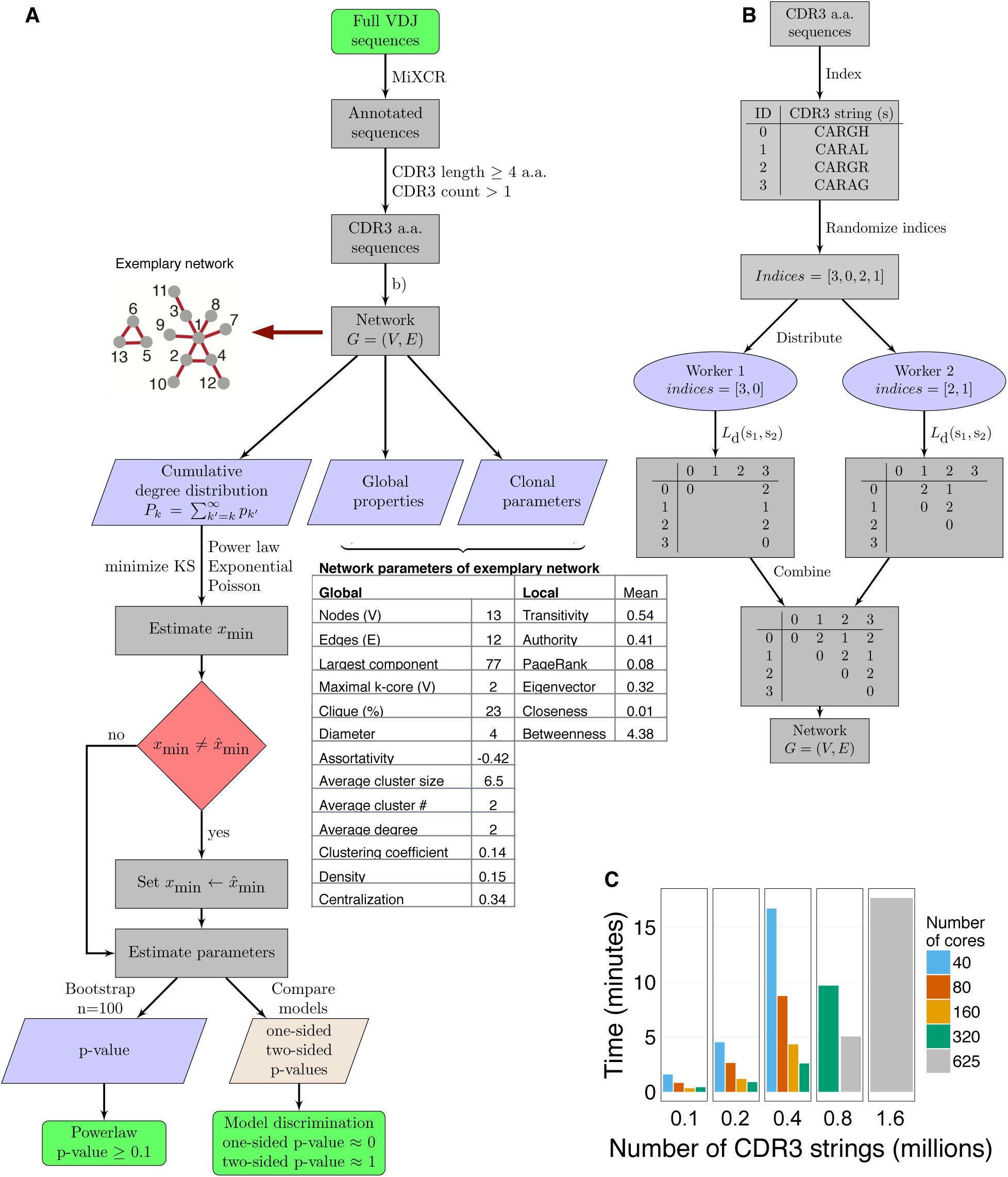
High-performance computing platform to construct and analyze large-scale networks from entire antibody repertoires. A. Data preprocessing, network construction and model fits to degree distribution (see Methods, Degree distribution fits for further details). Network parameters (global and mean local/clonal) are shown for the exemplary network.
B. Software schematics showing the distributed parallel computing platform used to partition the work among a cluster of many workers.
C. Computation time to construct large-scale networks depends on the number of CDR3 sequences and the number of cores used.

**Figure S2.**
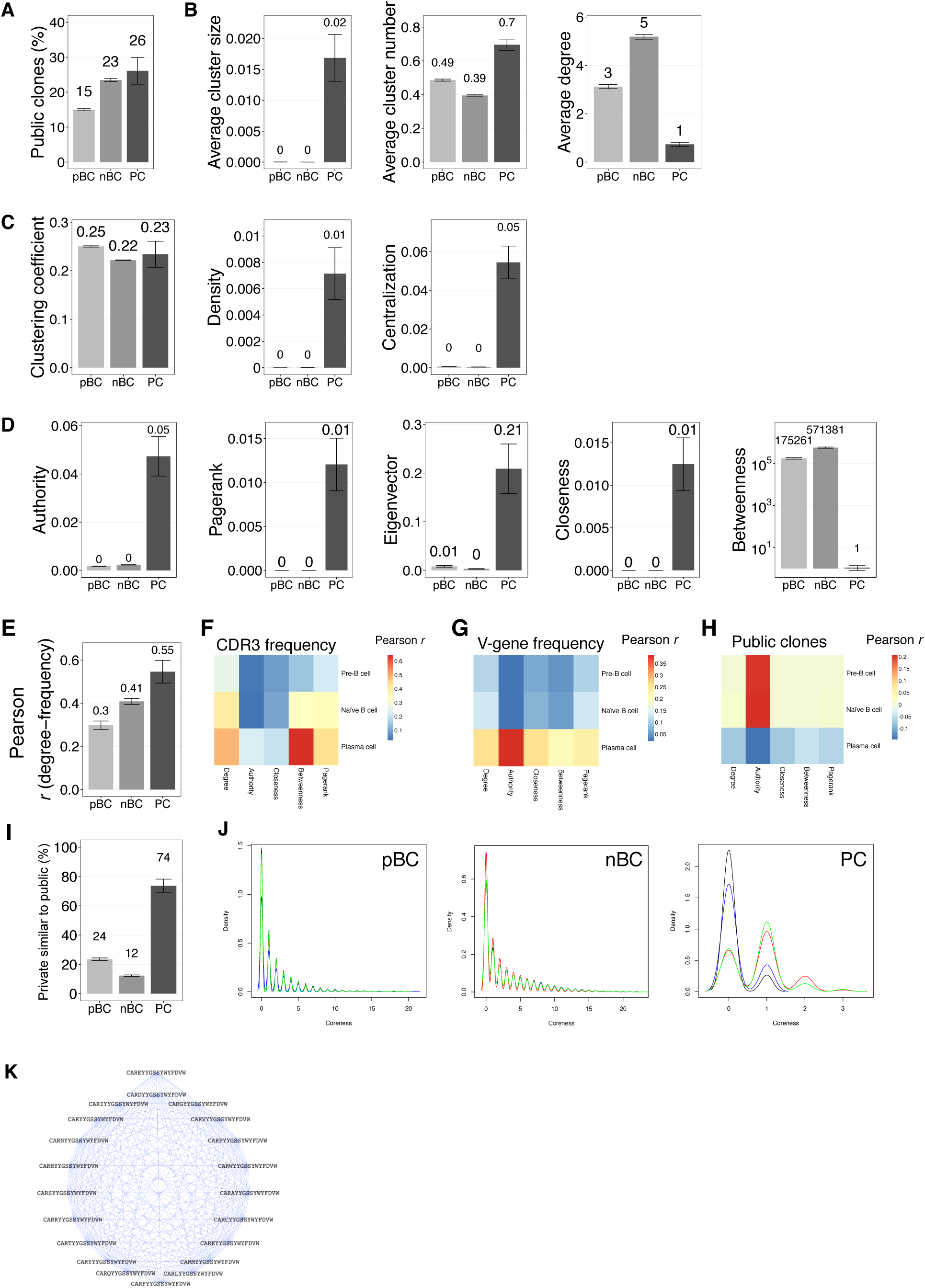
Global and clonal (local) network parameters of antibody repertoires of pre-B cells (pBC), naïve B cells (nBC) and plasma cells (PC). (A) Percentage of public clones, shared CDR3 clones between mice in pre-B cell (pBC), naïve B cell (nBC) and plasma cell (PC) repertoires.
(B–C) Global properties: Cluster analysis shows the average normalized cluster size and cluster number in the antibody repertoire networks. Average degree, clustering coefficient, density and (degree) centralization characterize the networks at the global level.
(D) Local properties: authority, PageRank, eigenvector, closeness and betweenness describe each clone in the network. Average values are shown for each B cell population, pre-B cells (pBC), naïve B cells (nBC) and plasma cells (PC). Barplots show mean±s.e.m, mice n=19.
(E) Pairwise Pearson correlation (*r*, mean±s.e.m) of CDR3 degree with CDR3 frequency in pre-B cells (pBC), naïve B cells (nBC) and plasma cells (PC) antibody repertoire networks.
(F) Pairwise Pearson correlation of local properties with CDR3 frequency (median, mice n=19).
(G) Pairwise Pearson correlation of local properties with germline V-gene frequency (mean, mice n=19).
(H) Pairwise Pearson correlation of CDR3 clonal (local) properties with public (1) vs. non-public (0) CDR3 clones (mean, mice n=19).
(I) Percentage of public clones similar (connected) to at least one other public CDR3 clone sequence by cohort (mean, mice n=19).
(J) Coreness density distribution for the unimmunized cohort of pre-B cells (pBC), naïve B cells (nBC) and plasma cells (PC). The x-axis shows the *k*-core (after removing sequentially shells of nodes of degree k-1). Line colors depict different mice.
(K) Example of CDR3 clones in the largest clique (complete connected subgraph) from a pre-B cell repertoire (NP-HEL cohort). For S2A-E, I, barplots represent mean±s.e.m; for each B-cell stage, n=19 mice.

**Figure S3.**
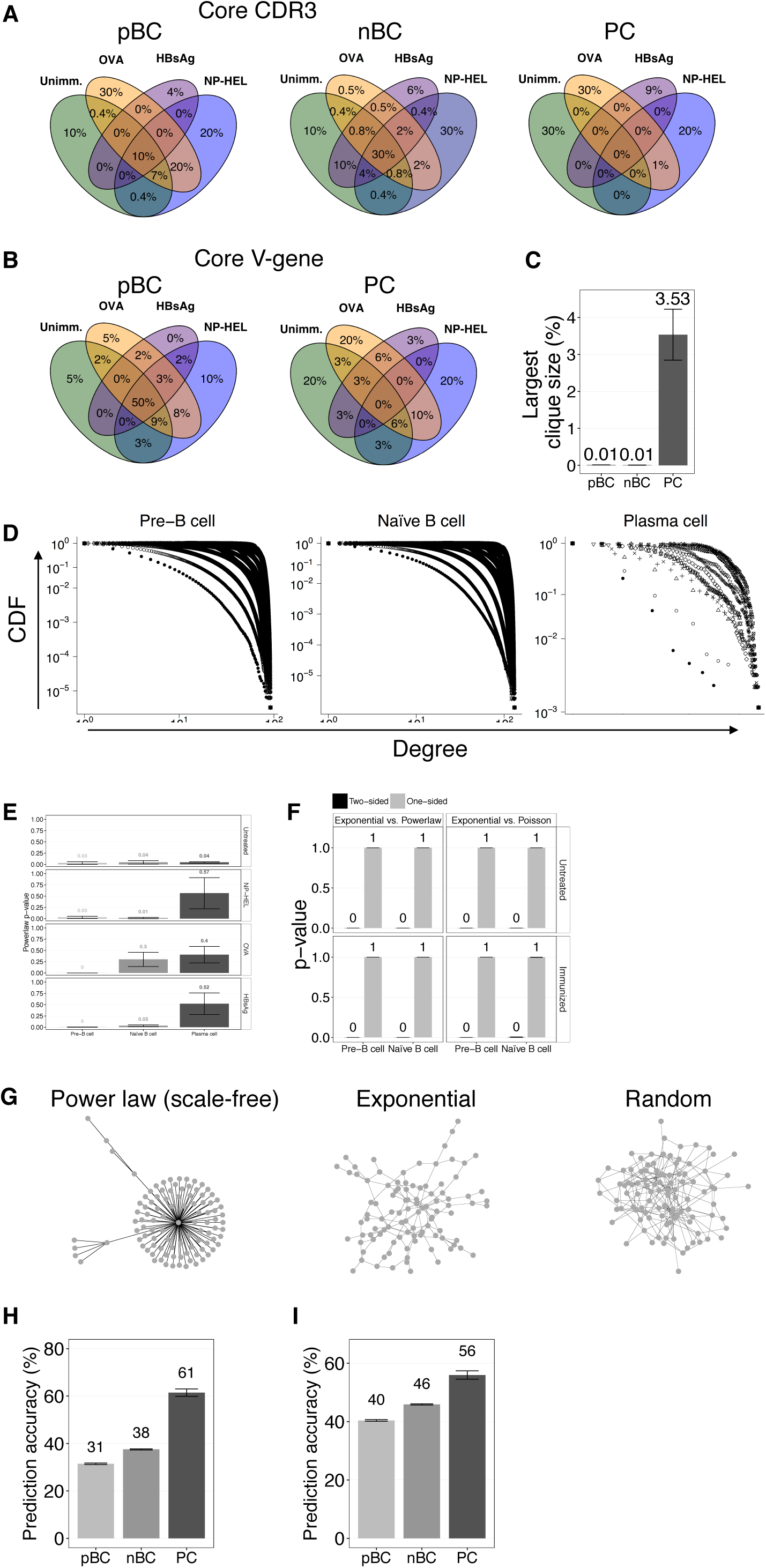
Core and structure (degree distributions of CDR3 similarity) analysis, and similarity layer prediction of antibody repertoire networks. A. Maximal core CDR3 clones overlap in pre-B cells (pBC), naïve B cells (nBC) and plasma cell (PC) repertoire networks.
B. Maximal core germline V-genes overlap in pre-B cells and plasma cell.
C. Percentage of the largest cliques (completely connected subgraph) along B cell development. Barplots represent mean±s.e.m; for each B-cell stage, n=19 mice.
D. Cumulative degree distributions (CDF). Each distribution line (different symbols) depicts one similarity layer LD_1–12_ (HBsAg-immunized mouse n. 4).
E. p-values (mean±s.e.m) of the power-law fit for each cohort.
F. One-sided and two-sided p-values (mean±s.e.m) for the discrimination between the exponential (one-sided p-value=1, two-sided p-value=0) and the power-law fits.
G. Graphics of power-law (γ =2.2), exponential and random network models of 100 nodes.
H. Prediction accuracy (*Q^2^*, leave-one-out cross-validated *R^2^*, mean±s.e.m) of selected distant similarity layers LD_4–12_ from LD_1_.
I. Prediction accuracy (*Q^2^*, mean±s.e.m) of all similarity layers (LD_2–12_) from LD_1_.

## REFERENCES

1. Bonsignori, M. et al. Antibody-virus co-evolution in HIV infection: paths for HIV vaccine development. Immunol. Rev. 275, 145–160 (2017).

2. Romero, P. et al. The Human Vaccines Project: A roadmap for cancer vaccine development. Sci. Transl. Med. 8, 334ps9–334ps9 (2016).

3. Hozumi, N. & Tonegawa, S. Evidence for somatic rearrangement of immunoglobulin genes coding for variable and constant regions. Proc. Natl. Acad. Sci. 73, 3628–3632 (1976).

4. Murphy, K., Travers, P. & Walport, M. Janeway’s immunobiology. N. Y. NY Garland Sci. (2012).

5. Weinstein, J. A., Jiang, N., White, R. A., Fisher, D. S. & Quake, S. R. High-Throughput Sequencing of the Zebrafish Antibody Repertoire. Science 324, 807–810 (2009).

6. Tonegawa, S. Somatic generation of antibody diversity. Nature 302, 575–581 (1983).

7. Bashford-Rogers, R. J. M. et al. Network properties derived from deep sequencing of human B-cell receptor repertoires delineate B-cell populations. Genome Res. 23, 1874–1884 (2013).

8. Ben-Hamo, R. & Efroni, S. The whole-organism heavy chain B cell repertoire from Zebrafish self-organizes into distinct network features. BMC Syst. Biol. 5, 27 (2011).

9. Chang, Y.-H. et al. Network Signatures of IgG Immune Repertoires in Hepatitis B Associated Chronic Infection and Vaccination Responses. Sci. Rep. 6, 26556 (2016).

10. Hoehn, K. B. et al. Dynamics of immunoglobulin sequence diversity in HIV-1 infected individuals. Philos. Trans. R. Soc. B Biol. Sci. 370, 20140241 (2015).

11. Lindner, C. et al. Diversification of memory B cells drives the continuous adaptation of secretory antibodies to gut microbiota. Nat. Immunol. 16, 880–888 (2015).

12. Klein, F. et al. Somatic Mutations of the Immunoglobulin Framework Are Generally Required for Broad and Potent HIV-1 Neutralization. Cell 153, 126–138 (2013).

13. Wu, X. et al. Maturation and Diversity of the VRC01-Antibody Lineage over 15 Years of Chronic HIV-1 Infection. Cell 161, 470–485 (2015).

14. Zaharia, M., Chowdhury, M., Franklin, M. J., Shenker, S. & Stoica, I. Spark: Cluster Computing with Working Sets. HotCloud 10, 95 (2010).

15. Lee, S. H., Kim, P.-J. & Jeong, H. Statistical properties of sampled networks. Phys. Rev. E 73, (2006).

16. Sethu, H. & Chu, X. A new algorithm for extracting a small representative subgraph from a very large graph. ArXiv Prepr. ArXiv12074825 (2012).

17. Amit, I. et al. Unbiased Reconstruction of a Mammalian Transcriptional Network Mediating Pathogen Responses. Science 326, 257–263 (2009).

18. Newman, M. Networks: an introduction. (Oxford University Press Inc., 2010).

19. Pavlopoulos, G. A. et al. Using graph theory to analyze biological networks. BioData Min. 4, 10 (2011).

20. Newman, M. E. Assortative mixing in networks. Phys. Rev. Lett. 89, 208701 (2002).

21. Freeman, L. C. Centrality in social networks conceptual clarification. Soc. Netw. 1, 215–239 (1978).

22. Bonacich, P. Power and centrality: A family of measures. Am. J. Sociol. 92, 1170–1182 (1987).

23. Brin, S. & Page, L. The anatomy of a large-scale hypertextual web search engine. Comput. Netw. ISDN Syst. 30, 107–117 (1998).

24. Barabási, A.-L. & Oltvai, Z. N. Network biology: understanding the cell’s functional organization. Nat. Rev. Genet. 5, 101–113 (2004).

25. Seidman, S. B. Network structure and minimum degree. Soc. Netw. 5, 269–287 (1983).

26. Galson, J. D. et al. In-Depth Assessment of Within-Individual and Inter-Individual Variation in the B Cell Receptor Repertoire. Front. Immunol. 6, (2015).

27. Horns, F. et al. Lineage tracing of human B cells reveals the in vivo landscape of human antibody class switching. eLife 5, e16578 (2016).

28. Jiang, N. et al. Lineage structure of the human antibody repertoire in response to influenza vaccination. Sci. Transl. Med. 5, 171ra19–171ra19 (2013).

29. Jackson, K. J. L., Kidd, M. J., Wang, Y. & Collins, A. M. The Shape of the Lymphocyte Receptor Repertoire: Lessons from the B Cell Receptor. Front. Immunol. 4, (2013).

30. Wagner, A. Robustness against mutations in genetic networks of yeast. Nat. Genet. 24, 355–361 (2000).

31. Burton, D. R. & Hangartner, L. Broadly Neutralizing Antibodies to HIV and Their Role in Vaccine Design. Annu. Rev. Immunol. 34, 635–659 (2016).

32. Briney, B. et al. Tailored Immunogens Direct Affinity Maturation toward HIV Neutralizing Antibodies. Cell 166, 1459–1470.e11 (2016).

33. Payne, J. L. & Wagner, A. The robustness and evolvability of transcription factor binding sites. Science 343, 875–877 (2014).

34. Burnet, S. F. M. The clonal selection theory of acquired immunity. 3, (Vanderbilt University Press Nashville, 1959).

35. Greiff, V., Miho, E., Menzel, U. & Reddy, S. T. Bioinformatic and Statistical Analysis of Adaptive Immune Repertoires. Trends Immunol. 36, 738–749 (2015).

36. Yaari, G. & Kleinstein, S. H. Practical guidelines for B-cell receptor repertoire sequencing analysis. Genome Med. 7, (2015).

37. Sidhu, S. S. & Fellouse, F. A. Synthetic therapeutic antibodies. Nat. Chem. Biol. 2, 682–688 (2006).

38. Logan, A. C. et al. High-throughput VDJ sequencing for quantification of minimal residual disease in chronic lymphocytic leukemia and immune reconstitution assessment. Proc. Natl. Acad. Sci. 108, 21194–21199 (2011).

39. Stern, J. N. et al. B cells populating the multiple sclerosis brain mature in the draining cervical lymph nodes. Sci. Transl. Med. 6, 248ra107–248ra107 (2014).

40. Jerne, N. K. Towards a network theory of the immune system. in Annales d’immunologie 125, 373–389 (1974).

41. Menzel, U. et al. Comprehensive Evaluation and Optimization of Amplicon Library Preparation Methods for High-Throughput Antibody Sequencing. PLoS ONE 9, e96727 (2014).

42. Bolotin, D. A. et al. MiXCR: software for comprehensive adaptive immunity profiling. Nat. Methods 12, 380–381 (2015).

43. Lefranc, M.-P. et al. IMGT, the international ImMunoGeneTics database. Nucleic Acids Res. 26, 297–303 (1998).

44. Dave, A. et al. Graphframes: an integrated api for mixing graph and relational queries. in Proceedings of the Fourth International Workshop on Graph Data Management Experiences and Systems 2 (ACM, 2016).

45. Schult, D. A. & Swart, P. Exploring network structure, dynamics, and function using NetworkX. in Proceedings of the 7th Python in Science Conferences (SciPy 2008) 2008, 11–16 (2008).

46. Peixoto, T. P. The graph-tool python library. figshare (2014).

47. Walt, S. van der, Colbert, S. C. & Varoquaux, G. The NumPy array: a structure for efficient numerical computation. Comput. Sci. Eng. 13, 22–30 (2011).

48. Clauset, A., Shalizi, C. R. & Newman, M. E. J. Power-Law Distributions in Empirical Data. SIAM Rev. 51, 661–703 (2009).

49. Gillespie, C. S. Fitting heavy tailed distributions: the poweRlaw package. J. Stat. Softw. 64, (2015).

50. Virkar, Y. & Clauset, A. Power-law distributions in binned empirical data. Ann. Appl. Stat. 8, 89–119 (2014).

51. Newman, M. E. The structure and function of complex networks. SIAM Rev. 45, 167–256 (2003).

52. Newman, M. E. Random graphs as models of networks. ArXiv Prepr. Cond-Mat0202208 (2002).

53. Kleinberg, J. M. Authoritative sources in a hyperlinked environment. J. ACM JACM 46, 604–632 (1999).

54. Csárdi, G. & Nepusz, T. The igraph library. http://igraph.org/ (2006).

55. Hyndman, R. J. & Khandakar, Y. Automatic time series for forecasting: the forecast package for R. (Monash University, Department of Econometrics and Business Statistics, 2008).

56. R Core Team. R: A Language and Environment for Statistical Computing. R Found. Stat. Comput. (2016).

57. Wickham, H. ggplot2: elegant graphics for data analysis. Springer N. Y. (2009).

58. Gaujoux, R. & Seoighe, C. A flexible R package for nonnegative matrix factorization. BMC Bioinformatics 11, 367 (2010).

59. Bastian, M., Heymann, S., Jacomy, M. & others. Gephi: an open source software for exploring and manipulating networks. ICWSM 8, 361–362 (2009).

